# COnstrained Reference frame diffusion TEnsor Correlation Spectroscopic (CORTECS) MRI: A practical framework for high-resolution diffusion tensor distribution imaging

**DOI:** 10.1101/2022.10.20.512879

**Authors:** Alexandru V. Avram, Kadharbatcha S. Saleem, Peter J. Basser

## Abstract

Diffusion MRI studies with resolutions of a few hundred micrometers have consistently shown that in the cortex water diffusion occurs preferentially along radial and tangential orientations with respect to the cortical surface, in agreement with histology. These dominant orientations do not change significantly even if the relative contributions from microscopic water pools to the net voxel signal vary across studies that use different diffusion times, b-values, TEs, and TRs. With this in mind, we propose a practical new framework for measuring non-parametric diffusion tensor distribution (DTD) MRI by constraining the microscopic diffusion tensors of the DTD to be diagonalized using the same orthonormal reference frame of the mesoscopic voxel. In each voxel, the constrained DTD (cDTD) is completely determined by the correlation spectrum of the microscopic principal diffusivities associated with the axes of the voxel reference frame. Consequently, all cDTDs are inherently limited to the domain of positive definite tensors and can be reconstructed efficiently with numerical methods for solving Inverse Laplace Transform problems.

Moreover, cDTDs can be measured using only data acquired with conventional single diffusion encoding, which can be obtained more efficiently than measurements with multiple diffusion encoding. In tissues with radial symmetry, such as the cortex, we can further constrain the cDTD to contain only cylindrically symmetric diffusion tensors and measure the 2D correlation spectra of radial and tangential diffusivities. To demonstrate this framework, we perform numerical simulations and analyze high-resolution dMRI data. We image 2D cDTDs in the cortex and derive marginal distributions of radial and tangential diffusivities, distributions of the microscopic fractional anisotropies and mean diffusivities, as well as their 2D correlation spectra to quantify the shape-size characteristics of the microscopic diffusion tensors. Signal components corresponding to specific bands in the measured correlation spectra show high specificity to cortical laminar structures observed with histology. Our framework drastically simplifies the measurement of non-parametric DTDs and may be applied retrospectively to analyze existing high-resolution dMRI data. Moreover, the framework provides a non-parametric generalization of DTI and subsumes existing diffusion signal representations and tissue models, enabling their harmonization, cross-validation, and optimization in specific clinical applications characterizing tissue changes.

## Introduction

By quantifying the microscopic motions of water molecules diffusion MRI (dMRI) provides a sensitive clinical tool to ! non-invasively probe the tissue structures at length scales (≈ 5μm) much smaller than the voxel size. In isotropic and anisotropic tissues, the dMRI signal at low diffusion sensitizations (b-values) can be described phenomenologically using diffusion tensor imaging (DTI) [Basser et al., 1994a,b]. In DTI, the diffusion signal attenuation in each voxel is modeled using a diffusion tensor, **D**, which has 6 degrees of freedom. The diffusion tensor can be decomposed or diagonalized in an orthogonal reference frame whose principal coordinate axes are characterized by the eigenvectors *ϵ*_**1**_, *ϵ*_**2**_, *ϵ*_**3**_. The normalized orthogonal unit vectors along the principal tensor axes represent 3 degrees of freedom of **D** that define its orientation with respect to the laboratory reference frame. The scalar principal diffusivities λ_1_, λ_2_, λ_3_ corresponding to these directions represent the other 3 degrees of freedom of **D** and determine the mean diffusivity and diffusion anisotropy. In general, **D** can be written as:

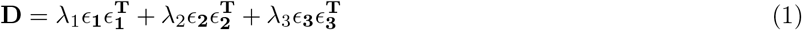

where *ϵ*_**1**_*ϵ*_1_^*T*^, *ϵ*_**2**_*ϵ*_**2**_^*T*^, *ϵ*_**3**_*ϵ*_**3**_^*T*^ are the principal coordinate axes dyads (or rank-1 tensors) derived from the eigenvectors of the diffusion tensor while the positivity of the principal diffusivities (i.e., eigenvalues of **D**) guarantees that **D** is positive definite.

However, at b-values larger than 1500*s*/*mm*^2^ the dMRI tissue signal is more sensitive to the intravoxel variation of water diffusion properties, and the DTI approximation may no longer hold. To quantify the intravoxel diffusion heterogeneity many approaches have been proposed, including using signal representations with higher-order terms, such as diffusion kurtosis imaging (DKI) [Jensen et al., 2005], generalized diffusion tensor imaging (GDTI) [Liu et al., 2004, Özarslan and Mareci, 2003], mean apparent propagator (MAP) MRI [Avram et al., 2016, Özarslan et al., 2013], as well as multi-exponential, multi-tensor, or multi-compartment tissue diffusion models [Assaf and Basser, 2005, Mulkern et al., 1999, Stanisz et al., 1997, Zhang et al., 2012].

Jian et al., extended the multi-tensor signal representations to describe intravoxel diffusion heterogeneity using a Wishart distribution of microscopic diffusion tensors [Jian et al., 2007]. Even though this parametric distribution is limited in its ability to accurately quantify the range of diffusion heterogeneity in healthy and diseased tissues, it nonetheless inspired great interest in measuring the underlying distribution of microscopic diffusion tensors (DTDs). In general, however, to disentangle microscopic processes with arbitrary diffusivities, diffusion anisotropies, and orientations, it is necessary to sensitize the measurement to diffusion-diffusion correlations [Callaghan and Komlosh, 2002, Cory et al., 1990, Mitra, 1995] by preparing the signal with multiple pulsed-field gradient (mPFG), or multiple diffusion encodings (MDE). Historically, biological and clinical applications of mPFG or MDE methods [Komlosh et al., 2007] have focused on estimating microstructural parameters such as the average axon diameters [Avram et al., 2013a,b, Koch and Finsterbusch, 2008, Komlosh et al., 2018] or pore size distributions [Benjamini et al., 2016]. More recently, MDE-prepared MRI measurements were described using tensor-valued diffusion encoding [Topgaard, 2017, Westin et al., 2016] in the context of probing diffusion heterogeneity in voxels composed of multiple non-exchanging Gaussian diffusion processes described with diffusion tensors whose corresponding ellipsoids have distinct sizes, shapes, and orientations, i.e., the DTD.

While, at least in principle, one can reconstruct DTDs from a very large number of measurements with encodings sampling the 6D space of b-tensors, in practice, the limited signal-to-noise ratio (SNR) and long scan duration make such clinical or biological experiments very challenging [Song et al., 2022, Topgaard, 2017]. To reduce the requirements for the high SNR level and a large number of measurement encodings some have made simplifying assumptions such as cylindrical symmetry of microscopic tensors [Topgaard, 2017] which reduce the dimensionality of non-parametric DTD reconstructions from six to four degrees of freedom. Alternatively, one can use parametric models (e.g., analytical functions) to estimate features of the DTDs [Jian et al., 2007, Magdoom et al., 2021, Szczepankiewicz et al., 2016, Westin et al., 2016] from data acquired using MDE and conventional single diffusion encoding (SDE) [Stejskal and Tanner, 1965].

Meanwhile, numerous studies using dMRI and other modalities provide converging evidence that, at a sufficiently small (i.e., mesoscopic) length scale, neuronal tissues, including cortical gray matter (GM) are organized preferentially along local orthogonal frames of reference. Ever since the earliest observations of cortical cyto- and myeloarchitecture [Brodmann, 1909, Cajal, 1909, Vogt, 1910], histochemistry and immunohistochemistry studies have consistently shown that cellular and subcellular structures at the microscopic scale are oriented predominantly along orthogonal, i.e., radial and tangential, orientations with respect to the cortical surface. This orthogonal reference frame persists at larger, mesoscopic scales of tens and hundreds of micrometers, and can be clearly seen in the arrangements of cells with various sizes, shapes and densities forming tissue architectural patterns along the same radial and tangential orientations such as cortical columns and laminae, respectively [Amunts and Zilles, 2015, Rubenstein and Rakic, 2020]. Most recently, studies using state-of-the-art electron microscopy (EM) in cortical GM [Lichtman and Denk, 2011, Shapson-Coe et al., 2021] have mapped the 3D organization of neuronal cells in gray matter with nanometer resolution over fields-of-view (FOVs) of hundreds of micrometers. These studies revealed in unprecedented detail anisotropic tissue structures, such as the microvasculature [Zhang et al., 2015], branching dendrites, neurofilaments, and other cell processes in various neuronal and non-neuronal cells (pyramidal neurons, intrinsic neurons, glial cells, etc.) roughly aligned along a local orthogonal frame of reference.

At mesoscopic length scales of a few hundred micrometers, diffusion processes in neural tissues align closely with the dominant orientations in the local tissue microstructure. Histological validation studies using ultra high-resolution dMRI have consistently found a good correspondence between the orientations of the underlying tissue microstructure and the orthogonal DTI reference frame [Budde and Annese, 2013, Seehaus et al., 2013, 2015] defined by *ϵ*_**1**_ϵ_**1**_^*T*^, *ϵ*_**2**_*ϵ*_**2**_^*T*^, *ϵ*_**3**_*ϵ*_**3**_^*T*^, or the fiber orientation distribution functions (FOD) [Tournier et al., 2004] measured with high-angular resolution diffusion MRI (HARDI) [Tuch et al., 2002] in the brain [Leergaard et al., 2010]. Numerous dMRI studies of cortical microstructure in fixed tissues [Aggarwal et al., 2015, Dyrby et al., 2011, Kleinnijenhuis et al., 2013, Leuze et al., 2014, McNab et al., 2009, 2013, Miller et al., 2011] and *in vivo* [Gulban et al., 2018, Heidemann et al., 2010, Jaermann et al., 2008, Kleinnijenhuis et al., 2015, McNab et al., 2013, Wang et al., 2021], for review see [Assaf, 2019], suggest that at submillimeter spatial resolution diffusion in the cortex is anisotropic and varies with the cortical folding geometry [Cottaar et al., 2018], in good agreement in with the cortical cyto- and myeloarchitectonic features observed with histology and other modalities [Nieuwenhuys, 2013]. Moreover, HARDI-derived FODs show preferentially radial and tangential components [Aggarwal et al., 2015, Kleinnijenhuis et al., 2013, Leuze et al., 2014] which evoke cortical columns [Petersen, 2007, Yacoub et al., 2008] and layers [Bastiani et al., 2016, Nagy et al., 2013], respectively, that can be observed with post-mortem histological staining. In addition, studies of laminar specific intra-cortical connectivity measured with diffusion fiber microtractography [Leuze et al., 2014] of cortical FODs [Aggarwal et al., 2015, Gulban et al., 2018] suggest a similar orthogonal (radial and tangential) organization.

Increasing the spatial resolution in dMRI reduces the intravoxel angular dispersion of subvoxel diffusion processes and implicitly the orientational variance of the DTD. At submillimeter spatial resolution, dMRI is sensitive to cortical diffusion anisotropy and allows us to identify the radial and tangential orientations along which diffusion processes align. Recently, a careful survey of the high-resolution dMRI literature [Assaf, 2019] suggests that when different contrast preparations are used to vary the relative contributions of microscopic tissue water pools to net voxel dMRI signal in the cortex, the dominant diffusion orientations, as measured using the DTI eigenvectors or the directions of FOD peaks, remain unaffected even though the relative diffusivities or FOD amplitudes along these orientations may change. At mesoscopic spatial resolutions of a few hundred micrometers, the orientational characteristics of the dMRI signal remain remarkably consistent across experiments with fixed and live cortical tissues using different T1- and/or T2-weightings, i.e., different echo time (TE), repetition time (TR), or inversion time (TI), diffusion sensitizations (b-values) or diffusion/mixing times. These findings imply that at mesoscopic spatial resolutions, subvoxel cortical diffusion tensors from microscopic water pools are coincident along the same dominant (radial and tangential) orientations and may have potentially different diffusion anisotropies and diffusivities. Implicitly, the DTD is predominantly determined by the variations in the shapes (diffusion anisotropies) and sizes (diffusivities) of the microscopic diffusion tensors, rather than by their relative orientations.

In this study, we describe a new framework that simplifies the measurement and analysis of diffusion heterogeneity in microscopic water pools within gray matter using a non-parametric DTD. Specifically, if the voxel size is small enough compared to the curvature of the cortex, we can constrain all the microscopic (subvoxel) diffusion tensors to share the same principal reference frame determined, for instance, by the dyadic of the principal diffusion eigenvectors, *ϵ*_**1**_, *ϵ*_**2**_, *ϵ*_**3**_, measured with DTI. With this constraint, the DTD is completely characterized by the voxel reference frame *ϵ*_**1**_*ϵ*_**1**_^*T*^,*ϵ*_**2**_*ϵ*_**2**_^*T*^,*ϵ*_**3**_*ϵ*_**3**_^*T*^, and by the 3D joint distribution of corresponding subvoxel principal diffusivities, λ_1_, λ_2_, λ_3_, which are random variables. This joint probability distribution can be estimated with a 3D Inverse Laplace Transform analysis using only single diffusion encoded (SDE) MR measurements. This practical, non-parametric framework for mapping DTDs, called COnstrained Reference frame diffusion TEnsor Spectroscopic (CORTECS) MRI, could quantify a wide range of cortical diffusion heterogeneity in healthy or diseased brains.

## Methods

### Higher spatial resolution reduces the intravoxel orientational dispersion

The net diffusion signal in an imaging voxel containing complex tissue microstructure can be described generally using an ensemble of subvoxel (i.e., microscopic) diffusion tensors with different sizes, shapes, and orientations, assumed to be in slow exchange, i.e., the diffusion tensor distribution (DTD). Ordinarily, we can quantify DTDs by analyzing diffusion-weighted images (DWIs) acquired with multidimensional diffusion encoding (MDE) [Magdoom et al., 2021, Topgaard, 2017, Westin et al., 2016]. The net dMRI voxel signal, *S*, is a function of the tensor-valued encoding variable called the b-tensor, **b**, computed by integrating the time-dependent diffusion gradient waveforms amplitudes, and is related to the underlying DTD, *p*(**D**):

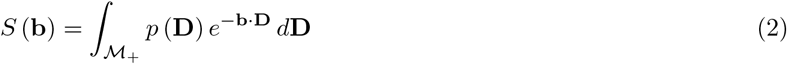

where the integral runs over the space or domain of all positive definite matrices, 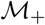. Since the random variable D has 6 degrees of freedom, *p*(**D**) is essentially a 6-dimensional joint probability distribution (or correlation spectrum) of the diffusion tensor elements. The high dimensionality and the inherent challenge of defining the subspace of positive-definite random tensor-valued variables, **D**, make solving this problem infeasible in practice, as no closed-form solution exists. Measuring *p*(**D**) requires a prohibitively large number of measurements with a very high signal-to-noise ratio (SNR) and MDE. Previously, approximations to *p*(**D**) have been proposed either by assuming parametric models and/or by using statistical reconstruction algorithms [Jian et al., 2007, Magdoom et al., 2021, Szczepankiewicz et al., 2016, Topgaard, 2017, Westin et al., 2016].

In cortical GM the orthogonal coordinate axes along which diffusive fluxes align at the microscopic scale of cellular and subcellular structures (i.e., diffusion length scale) are propagated at larger mesoscopic scales guiding the assembly of these structures into orthogonal tissue architectural patterns of cortical laminae and columns [Nieuwenhuys, 2013, Rubenstein and Rakic, 2020]. If the voxel size of dMRI data is significantly smaller than the minimum radius of the curvature of the underlying anatomy (i.e., cortical folding) the orientational variance of subvoxel (microscopic) diffusion processes can be neglected (Fig. 1). Microscopic diffusion processes are coincident with the axes of the local microstructural reference frame determined by the cortical cyto- and myeloarchitecture. For a continuously varying cortical anatomy with a minimum radius of curvature, *R*, the range of orientational misalignment between the microscopic diffusion tensors and the voxel reference frame, ±*θ_max_*, in a cubic voxel of side length, *x*, is:

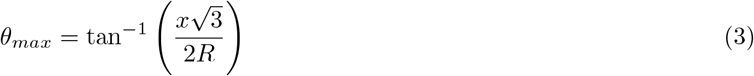

**Figure 1.**
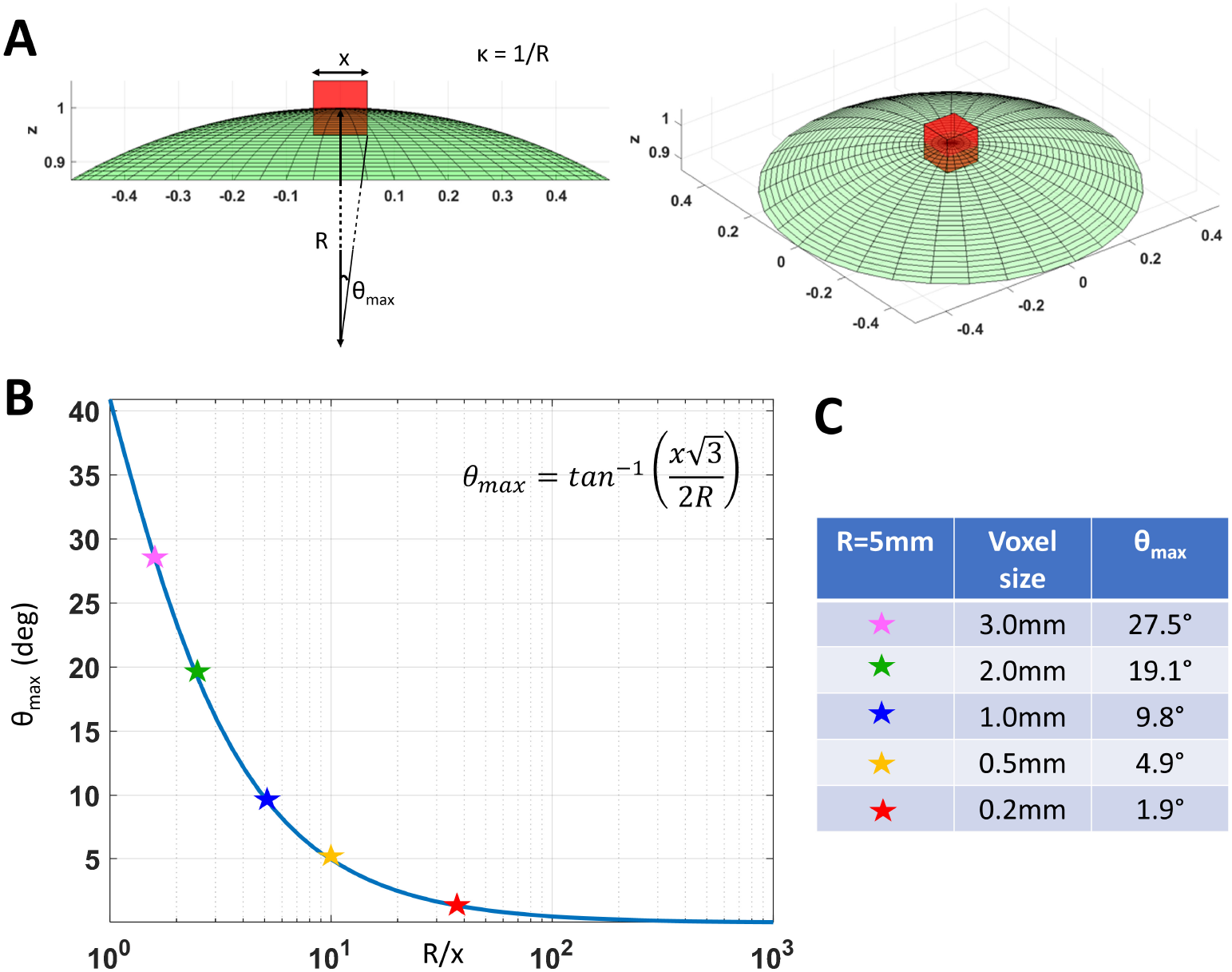
**A.** As we decrease the voxel size, *x*, relative to the radius of curvature of the tissue (e.g., due to cortical folding), *R*, the intravoxel orientational variance of the continuously varying microstructural reference frame also decreases. For a voxel with an arbitrary orientation relative to the underlying microstructure, the range of intravoxel orientational variation due to tissue curvature is ±*θ_max_*. **B.** The value of *θ_max_* decreases rapidly at low spatial resolutions, *R/x*, but changes very slowly at higher spatial resolutions, *R/x*. **C.** A quantitative comparison of *θ_max_* at different voxel sizes assuming a cortical radius of curvature *R* = 5*mm* shows the significant reduction in intravoxel orientational variance due to the effects of anatomical curvature at high spatial resolutions.

Fig. 1B shows that *θ_max_* decreases rapidly at low spatial resolutions, 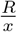, but changes slowly at higher values of 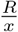 (Fig. 1B). At a spatial resolution of a few hundred micrometers the voxel size is much smaller than the cortical radius of curvature (R=5mm) leading to very small values of *θ_max_*. Under these circumstances, it is reasonable and practical to constrain all diffusion tensor processes in microscopic water pools throughout the voxel (i.e., the DTD) to be described using the same local orthogonal reference frame.

### COnstrained Reference frame diffusion TEnsor Correlation Spectroscopic (CORTECS) MRI

Fixing the local reference frame for all subvoxel tensors has several surprising advantages. First, it significantly reduces the dimensionality of *p*(**D**) and decouples the statistical random variables needed to describe *p*(**D**). Specifically, the 6D vector/tensor random variable, **D**, corresponding to the 6 components (or degrees of freedom) needed to describe the general DTD is reduced to a 3D random variable comprising the three principal diffusivities, λ_1_, λ_2_, λ_3_ along the axes of the fixed voxel frame of reference, *ϵ*_**1**_*ϵ*_**1**_^*T*^, *ϵ*_**2**_*ϵ*_**2**_^*T*^, *ϵ*_**3**_*ϵ*_**3**_^*T*^, respectively, which are sufficient to describe the constrained DTDs (cDTDs) within the Coordinate Reference frame diffusion Tensor Correlation Spectroscopic (CORTECS) MRI framework (Fig. 2A,B). Using the eigenvalue decomposition of the diffusion tensor (Eq. 1) we can re-write Eq. 2 as a more tractable 3D Inverse Laplace transform (ILT) problem:

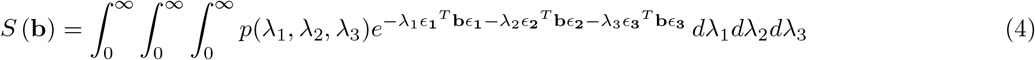

where *ϵ*_**i**_^*T*^**b***ϵ*_*i*_ is a non-negative scalar weighting (quadratic form) that represents the reciprocal Laplace variable corresponding to λ_*i*_. Besides the drastic reduction in the computational complexity due to the dimensionality reduction, the CORTECS framework inherently enforces positive definiteness of diffusion tensors by requiring positivity of the λ_*i*_.

**Figure 2.**
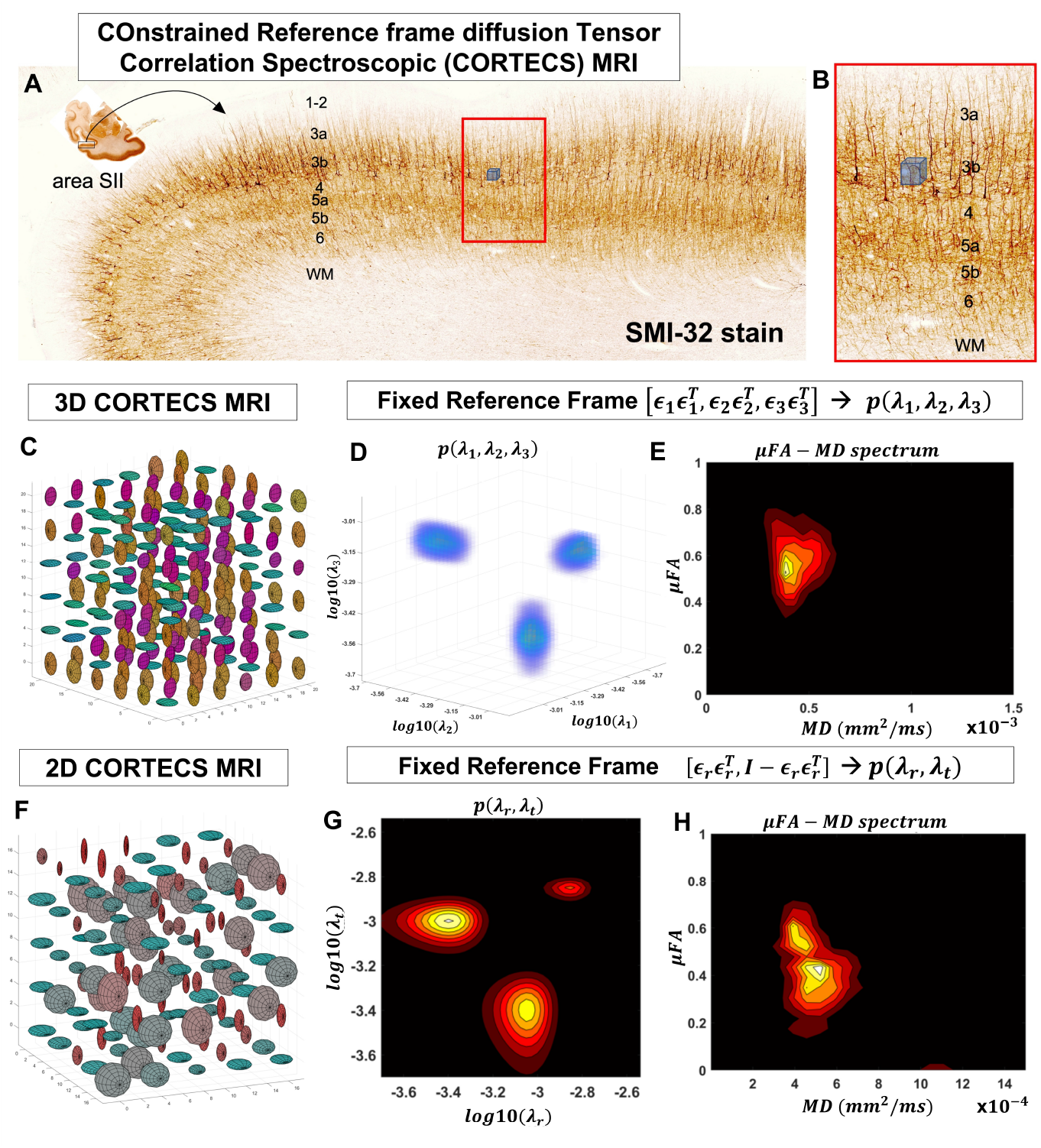
At a mesoscopic length scale cortical cyto- and myeloarchitecture is organized preferentially along the axes of an orthogonal frame of reference (**A**). If the dMRI spatial resolution is sufficiently small (**Fig. 1**) we can measure DTDs efficiently using the constraints of the CORTECS MRI framework (**B**). If we constrain all microscopic diffusion tensors to have the same principal axes of diffusion (**C**) we can quantify the DTD as the 3D correlation spectrum of the corresponding principal diffusivities (**D**). If the microarchitecture varies along a single radial orientation we can further constrain the DTD to contain only axisymmetric tensors (**F**) and quantify the 2D correlation spectrum of the corresponding radial and tangential diffusivities. We can also quantify the shape-size (i.e., microscopic FA-MD) correlation spectra of microscopic tensors from the 3D (**E**) or 2D (**H**) constrained reference frame DTDs (cDTDs).

Another very important advantage of constraining the reference frames of the DTD tensor random variable is that we can measure *p*(λ_1_, λ_2_, λ_3_) using only DWIs with single pulse-field gradient (sPFG) or single diffusion encoding (SDE), a.k.a. linear tensor encoding with rank-1 b-tensors. For a conventional SDE DWI with an arbitrary b-value, *b*, and diffusion gradient direction given by the unit vector **g** = [*g_x_,g_y_,g_z_*]^*T*^, the encoding b-tensor has rank-1, **b** = *b***gg**^*T*^. We can rewrite the signal equation above with respect to the components of **g** expressed in the voxel frame of reference, **g**’ = [*g*’_1_, *g*’_2_, *g*’_3_]^*T*^ = [*ϵ*_**1**_*ϵ*_**2**_*ϵ*_**3**_] **g**:

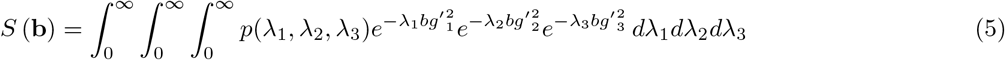

The factors 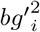 are the non-negative weighting parameters of the principal diffusivities, λ_*i*_, in the Laplace Transform representation of the signal. We can generate a wide range of joint weighting parameters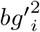 by varying the b-value and diffusion gradient orientations in conventional SDE preparations. Subsequently, from multiple SDE DWIs we can estimate, in each voxel, the correlation spectrum of principal diffusivities, *p*(λ_1_, λ_2_, λ_3_) which quantifies the properties of all microscopic diffusion tensor processes. Compared to MDE-DWIs, the conventional SDE-DWI can be acquired efficiently using product single pulsed-field gradient (sPFG) spin-echo (SE) diffusion MR sequences [Stejskal and Tanner, 1965] available on all microimaging and clinical MRI scanners. In general, SDE-DWIs can achieve higher b-values, shorter echo times (TEs), higher spatial resolution, and/or better SNR than MDE-DWIs using double or triple diffusion encoding. Moreover, the spectral reconstruction of *p*(λ_1_, λ_2_, λ_3_), henceforth referred to as 3D cDTD, does not require statistical methods to enforce positive definiteness but can still benefit from various techniques that may be used to solve ILT-like problems, such as L_2_- or L_1_-norm regularization, compressed sensing [Bai et al., 2015], or constrained optimization [Benjamini et al., 2016], etc.

If the underlying microstructure is radially symmetric, i.e., varying along a single preferred orientation, we can make an additional simplification to the problem and assume locally oriented cylindrical symmetry for each ensemble of subvoxel diffusion tensors (Fig. 2F,G,H). In this case, the voxel reference frame is determined by a single orientation, *ϵ*_**1**_*ϵ*_**1**_^*T*^, i.e., the radial direction, which implicitly defines the orthogonal, tangential component described by the rank-2 tensor 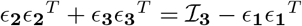, where 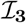 is the 3×3 identity matrix. We can relate the signal in a voxel with fixed principal axis *ϵ*_**1**_*ϵ*_**1**_^*T*^ to a two-dimensional correlation spectrum of radial and tangential diffusivities, *p*(λ_*r*_, λ_*t*_) that completely determines the corresponding cylindrically symmetric DTD:

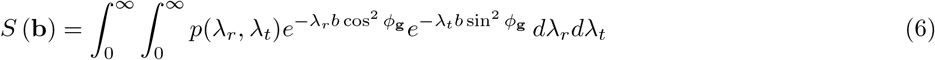

The parameter *ϕ*_**g**_ = arccos(*ϵ*_**1**_^*T*^_**g**_) represents the angle between the applied gradient direction, **g**, and the radial direction of the underlying reference frame, *ϵ***1***ϵ*_**1**_^*T*^. In radially symmetric tissues such as the cortex with cytoarchitecture aligned along a dominant radial direction (columns) and the corresponding tangential plane (laminae), diffusion processes are likely oriented and cylindrically symmetric and can be more economically and effectively quantified using this lower-dimensional correlation spectrum, called 2D cDTD.

Lastly, in a final simplifying step, if all subvoxel diffusion processes are isotropic, the correlation spectrum of diffusion tensor eigenvalues reduces to a distribution of a single scalar diffusivity random variable, λ_0_, which can be viewed as 1D cDTD:

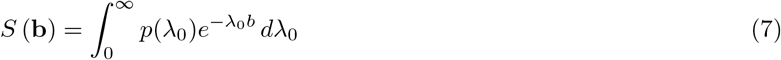

As an aside, we should point out an important connection between 1D cDTD MRI and our previously proposed methods for one- and multidimensional MD spectroscopic MRI using isotropic diffusion encoding (IDE) [Avram et al., 2019, 2021]. Mapping non-parametric spectra of MD values in microscopic tissue water pools using multiple IDE measurements does not require that diffusion in these pools is isotropic. Meanwhile, the 1D cDTD MRI spectral reconstruction using Eq. 7 correctly quantifies the spectra of water mobilities only if all diffusion processes within the voxel are isotropic, in which case the two methods will provide congruent results.

### Mapping distributions and correlation spectra of microscopic fractional anisotropy and mean diffusivity

From the measured cDTD within each voxel, we can compute non-parametric distributions and correlation spectra of DTI-derived parameters of microscopic diffusion tensors, such as fractional anisotropy (FA) or mean diffusivity (MD). Specifically, we can define a new random variable, α, that quantifies the FA of each microscopic diffusion tensor in the cDTD:

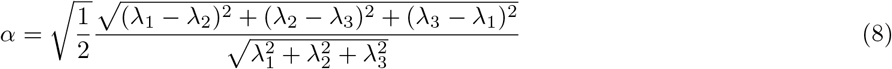

From *p*(λ_1_, λ_2_, λ_3_) we can then derive the probability density function (one-dimensional spectrum) of the microscopic tensor FAs, *p*_*FA*_(*α*), which quantifies the cDTD shape heterogeneity non-parametrically. The statistical moments of the *p_FA_*(*α*) provide important microstructural parameters, such as the microscopic anisotropy, *μFA*, computed as the mean of *p_FA_*(*α*). Similarly, we can define a new cDTD-derived random variable that quantifies the mean diffusivity of each microscopic tensor, *μ* = (λ_1_ + λ_2_ + λ_3_)/3, and compute the probability density function *P_MD_* (*μ*) to describe the spectrum the microscopic water mobilities in tissue non-parametrically.

Finally, from *p*(λ_1_, λ_2_, λ_3_) we can also compute non-parametric multidimensional correlation spectra of two or more microscopic DTI metrics. For example, we can quantify non-parametrically the correlations between the shapes and sizes of the diffusion ellipsoids corresponding to the underlying diffusion tensors by computing the joint probability density function of the two random variables *α* and *μ*, *p_FA-MD_* (*α, μ*). This practical and efficient decomposition of tissue heterogeneity based on diffusion anisotropy and mean diffusivity correlations in microscopic water pools may reveal specific microstructural motifs or patterns potentially relevant to many clinical applications.

### A generalization of various diffusion tensor signal models

The CORTECS framework can describe a wide range of heterogeneous diffusion processes in healthy and diseased tissues and subsumes several diffusion tensor signal models. For example, if we constrain 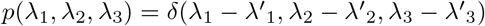, 3D cDTD simplifies to conventional DTI with the three mean eigenvalues 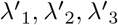. In this way, 3D cDTD can be viewed as a generalization of high-resolution DTI that quantifies intravoxel diffusion heterogeneity as a non-parametric correlation spectrum of the principal diffusivities in microscopic water pools. To describe multi-exponential or multi-tensor signal decays in heterogeneous tissues [Avram et al., 2020, Mulkern et al., 1999, Stanisz et al., 1997] we can assume that *p*(λ_1_, λ_2_, λ_3_) can be represented as a sum of delta functions (point masses) [Avram et al., 2020]. Moreover, the spectroscopic decomposition of the net voxel signal in cDTD makes it easy to disentangle partial volume contributions, such as those from cerebrospinal fluid (CSF), or free water in tissues caused by edema or other processes [Pasternak et al., 2009].

### Monte Carlo Simulations

We conducted Monte Carlo (MC) simulations to evaluate the numerical stability and accuracy of the voxel-wise estimation of 3D and 2D cDTDs from noisy data. Specifically, starting from ground truth DTDs constrained with fixed voxel reference frames (2D and 3D cDTDs), defined analytically using multidimensional lognormal distributions, respectively, we computed the dMRI signals expected from an experiment using conventional single-diffusion encoded (SDE) DWI measurements with the same gradient orientations and b-values as in our fixed-brain experiment described below. Next, from these ground truth signals, we generated 500 instances of noisy measurements by adding Rician noise to simulate real measurements with different SNR levels. From each set of noisy measurements, we computed the corresponding normalized 3D correlation spectra of principal diffusivities, or normalized 2D correlation spectra of radial and tangential diffusivities and compared the statistics of these spectra (mean and standard deviation) to the corresponding ground truth 3D and 2D DTDs, respectively.

### Ultra high-resolution dMRI of a fixed macaque monkey brain

The brain of a healthy young adult rhesus macaque monkey (*Macaca mulatta*) weighing 13.55 kg was prepared using a well-controlled perfusion fixation process, as described in [Saleem et al., 2021]. In brief, the animal was deeply anesthetized with sodium pentobarbital and perfused transcardially with heparinized saline, followed by 4% paraformaldehyde in 0.1 M phosphate buffer (pH 7.4). After perfusion, the brain was removed from the cranium and post-fixed for 8h in the same buffered paraformaldehyde solution. Following the post-fixation, the brain was transferred into 0.1 M phosphate-buffered saline with sodium azide before the MRI data acquisition. All procedures were carried out under a protocol approved by the Institutional Animal Care and Use Committee of the National Institute of Mental Health (NIMH) and the National Institute of Health (NIH) and adhered to the Guide for the Care and Use of Laboratory Animals (National Research Council).

Based on a preliminary structural MRI scan of the specimen, we fabricated a three-dimensional (3D) brain mold inside a cylindrical acrylic plastic container. The specimen was positioned inside the brain mold which was placed inside a custom 70mm cylindrical container. The container was filled with Fomblin and gently stirred under a vacuum for 4 hours to remove air bubbles. Subsequently, the container was sealed and prepared for MR imaging using a Bruker 7T horizontal-bore MRI scanner and a Bruker 72mm quadrature RF coil.

We acquired whole-brain diffusion-weighted images (DWIs) with a cubic voxel size of 200μm, i.e., a 375×320×230 imaging matrix on a 7.5×6.4×4.6cm field-of-view (FOV), using a diffusion spin-echo 3D echo-planar imaging (EPI) sequence with 50ms echo time (TE), 650ms repetition time (TR), 18 segments and 1.33 partial Fourier acceleration. We obtained a total of 112 DWIs using multiple b-value shells (100, 1000, 2500, 4500, 7000, and 10000 *s/mm*^2^) with diffusion-encoding gradient orientations (3, 9, 15, 21, 28, and 36, respectively) uniformly sampling the unit sphere on each b-value shell and across shells. The diffusion gradient pulse durations and separations were δ=6ms and △=28ms, respectively. Each DWI volume was acquired using a single average in 52 minutes. The total duration of the diffusion MRI scan was 93 hours and 20 minutes. We processed all whole-brain high-resolution DWIs with the TORTOISE software pipeline [Pierpaoli et al., 2010] which includes image registration, Gibbs ringing correction [Kellner et al., 2016], denoising [Veraart et al., 2016], corrections for EPI distortion including eddy currents and B0 inhomogeneities using a high-tissue contrast structural magnetization transfer (MT) scan as an anatomical template.

### Histological processing

After imaging, the perfusion-fixed brain specimen was prepared for histological processing with five different stains as described in [Saleem et al., 2021]. In brief, the brain blocks were frozen and serially sectioned through the entire brain at 50μm thickness in the coronal plane. Next, five sets of interleaved sections were processed for Parvalbumin (PV), neurofilament protein (SMI-32), choline acetyltransferase (ChAT), cresyl violet (CV), and Acetylcholinesterase (AchE) staining. Finally, we captured high-resolution images of stained sections using a Zeiss high-resolution slide scanner with a 20X objective. These images were then manually aligned with the corresponding slices from the MRI data for comparison of cortical architectonic features.

### 2D CORTECS MRI in the fixed macaque brain

From the distortion-corrected DWIs we estimated fiber orientation distribution functions and compared their orientations in the cortex to those of microscopic structures observed on histological images. We further analyzed the high-resolution DWIs using DTI and estimated the voxel reference frame, *ϵ*_**1**_*ϵ*_**1**_^*T*^, *ϵ*_**2**_*ϵ*_**2**_^*T*^, *ϵ*_**3**_*ϵ*_**3**_^*T*^, through eigenvalue decomposition of the net diffusion tensor in each voxel (Eq. 1). Subsequently, using the diffusion principal diffusion direction *ϵ*_**1**_*ϵ*_**1**_^*T*^, we computed the diffusion weightings of radial and tangential processes, *b* cos^2^ *ϕ*_**g**_ and *b* sin^2^ *ϕ*_**g**_, respectively, for each measurement encoding and in each voxel. Finally, we estimated a piecewise continuous approximation of the 2D cDTD correlation spectrum, *p*(λ_*r*_,λ_*t*_), by numerically solving the 2D ILT problem (Eq. 6) using linear least-squares error minimization with L2-norm regularization [Hansen, 1992] and positivity constraints. A detailed description of the implementation of the spectral reconstruction algorithm can be found in [Avram et al., 2019, 2021]. The spectral bins of the cDTDs reconstruction were defined on a 12 x 12 grid of logarithmically spaced λ_*r*_ and λ_*t*_ values ranging from 0.01 – 2.00*μm*^2^/*ms*. From the 2D cDTD correlation spectrum *p*(λ_*r*_,λ_*t*_) we derived maps of the marginal distributions of the radial and tangential diffusivities, microscopic FA and MD, as well as the microscopic FA-MD correlation spectra, *p_FA-MD_* (*α,μ*), and related these results to cortical cytoarchitectonic features observed with histology. The microscopic FA-MD correlation spectra were estimated numerically from the cDTDs using an 11 x 11 grid of microscopic FA and MD values. We empirically selected several *ad hoc* spectral domains in the 2D joint distributions *p*(λ_*r*_, λ_*t*_) and *p_FA-MD_*(*α, μ*) to best capture the most prominent spatial-spectral correlations. We compared maps of the signal components corresponding to these domains to the cortical cytoarchitectonic features in the corresponding stained tissue section. The cDTD reconstruction and analysis for the numerical simulations and fixed brain experiments were implemented in MATLAB.

## Results

### Monte Carlo Simulations

Monte Carlo (MC) simulations of 3D and 2D cDTD reconstructions show that it is possible to distinguish subvoxel diffusion tensor processes that are aligned in the same voxel reference frame based on differences in the correlations of their principal diffusivities using experimental designs that contain only SDE measurements and can be achieved with current MRI scanners. Fig. 3 shows the MC results for a ground truth 3D cDTD (i.e., correlation spectrum of principal diffusivities) that consists of a mixture of three multivariate log-normal distributions, reflecting the presence of 3 microscopic water pools with distinct diffusion tensor properties. The mean normalized spectra reconstructed from noisy measurements with various SNR levels provide good estimates for the locations and concentrations (i.e., areas under the peaks) of individual signal components. Meanwhile, at higher SNR levels, the exact shapes of the estimated spectral peaks are more accurately resolved. Lower dimensional marginal distributions derived from the 3D cDTDs also reveal the presence of multiple peaks and show improved accuracy at higher SNR levels. Similar results were obtained in MC simulations using 2D cDTDs shown in Fig. 4. The ground truth correlation spectrum of radial and tangential diffusivities, *p*(λ_*r*_, λ_*t*_), that defines the 2D cDTD consists of a mixture three microscopic diffusion processes described by mixtures of multivariate log-normal distributions. The locations and concentrations of these peaks can be estimated over a wide range of SNR levels, with improved accuracy at higher SNRs. Errors in the estimated spectra may be due to measurement noise, the limited number of measurements, and/or the regularization and positivity constraints used to improve the condition number of the spectral reconstruction.

**Figure 3.**
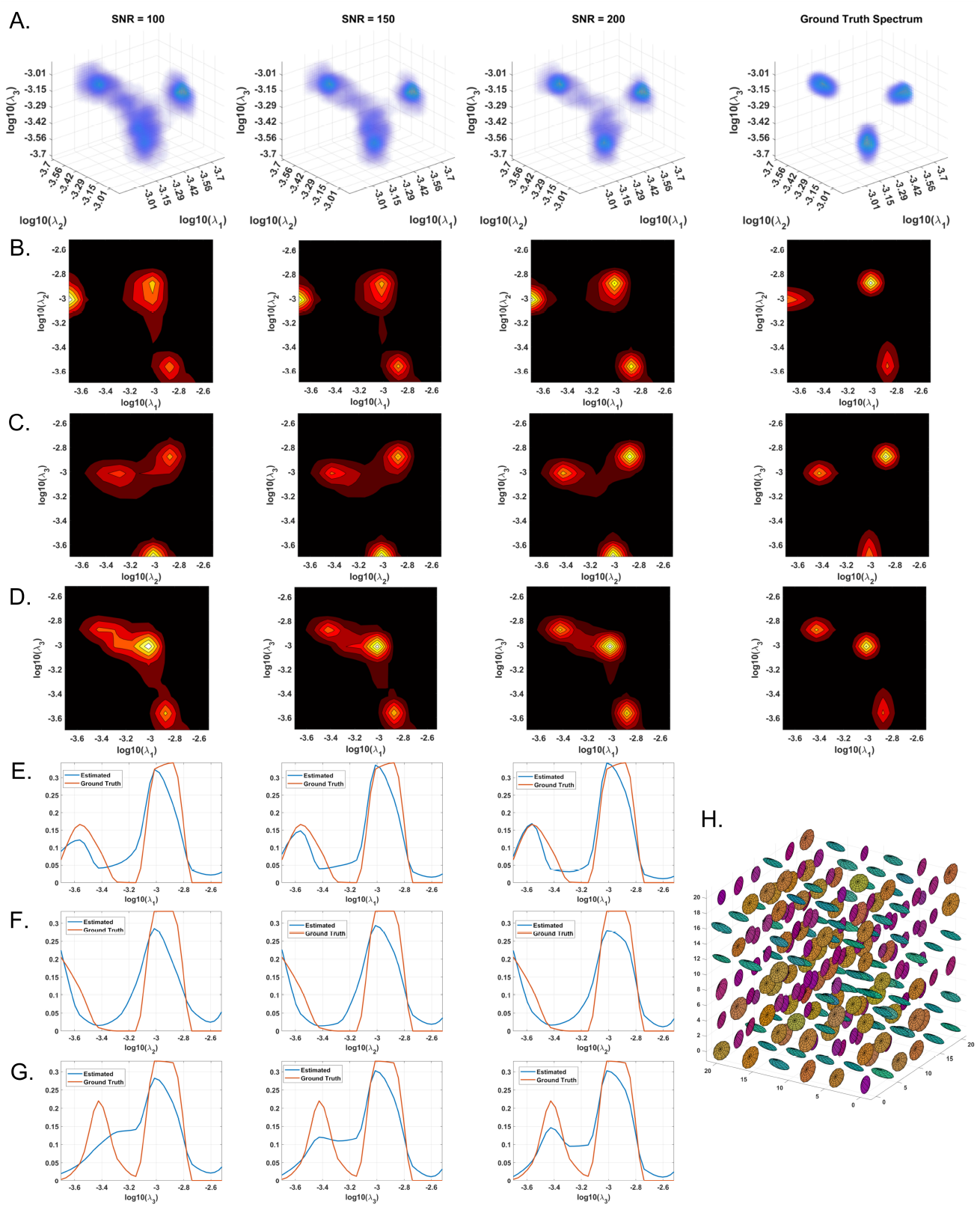
Monte-Carlo simulation results illustrating the accuracy and numerical stability of the 3D cDTD reconstruction as a function of SNR. **A**: Log-log-log plots of mean normalized 3D cDTD correlation spectra of the principal diffusivities reconstructed from data with different SNRs. **B,C,D**: Log-log plots of mean normalized 2D marginal distributions derived from the 3D cDTDs in the top row. **E,F,G**: Log plots of the mean normalized 1D marginal distributions derived from the 3D cDTDs in the top row. **H**: A numerically simulated illustration of an ensemble of diffusion tensors described by the ground truth 3D cDTD.

**Figure 4.**
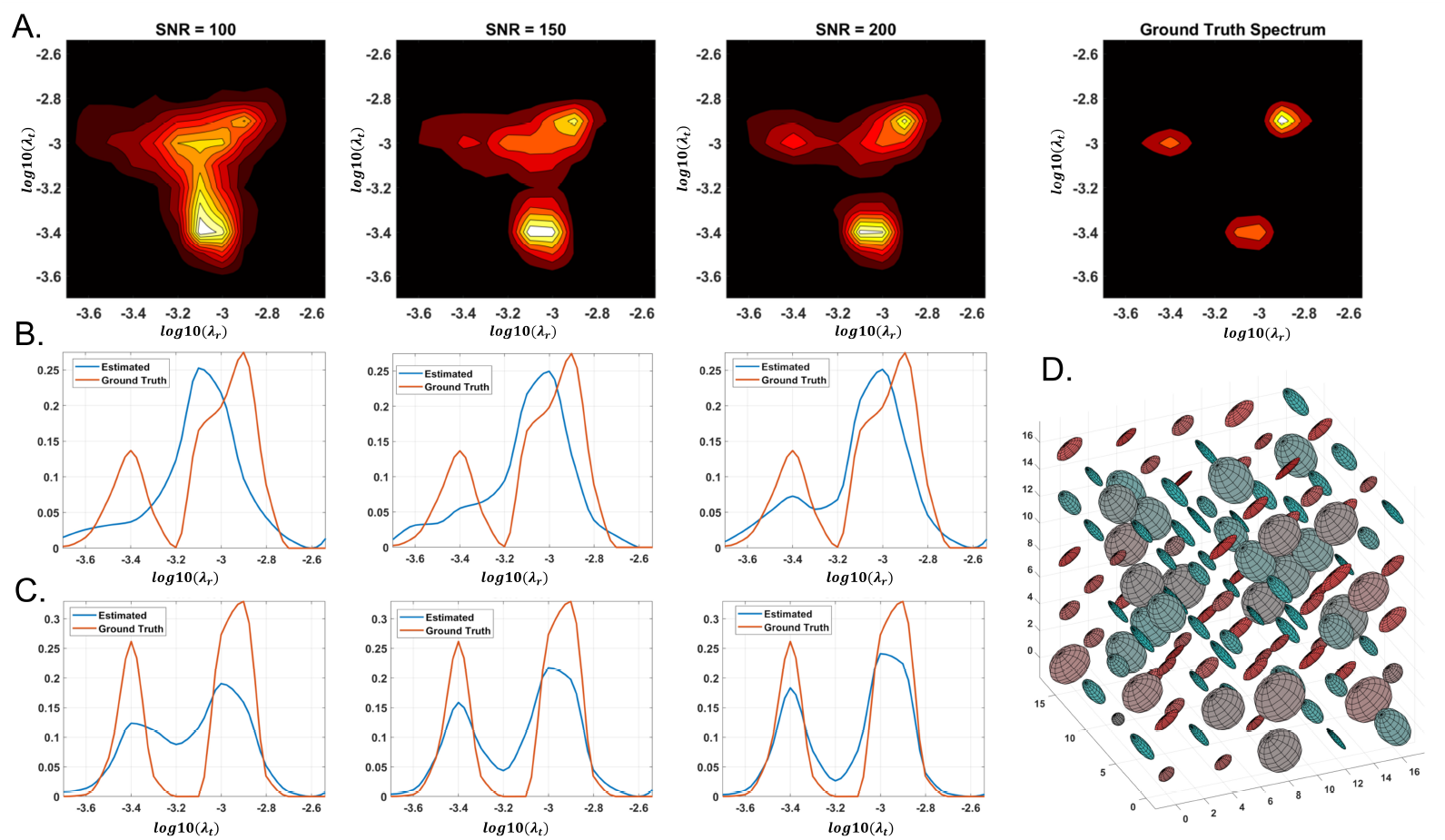
Monte-Carlo simulation results illustrating the accuracy and numerical stability of the 2D cDTD reconstruction as a function of SNR. **A**: Log-log plots of mean normalized 2D cDTD correlation spectra of principal diffusivities reconstructed at different SNR levels. **B,C**: Log plots of mean normalized 1D marginal distributions derived from the 2D cDTDs in the top row. **D**: A numerically simulated illustration of an ensemble of diffusion tensors described by the ground truth 2D cDTD.

The spectral resolution depends on the number of measurements with different encodings, the SNR level, and the use of constraints and regularization for spectral reconstruction. For a fixed SNR and a wide range of signal weightings (e.g., b-values), slowly decaying components have a better contrast-to-noise ratio (CNR) than fast decaying ones and can therefore be resolved with higher spectral resolution. The resulting non-uniform spectral resolution is not unique to CORTECS MRI but is inherent to the data required by all multidimensional relaxation and diffusion spectroscopic MRI methods. These techniques aim to disentangle multiexponential processes by quantifying the underlying distribution of decay constants non-parametrically using an ILT-like reconstruction from a finite set of measurements. The spectral resolution could be improved using more advanced spectral reconstruction algorithms that rely on statistical methods [Prange and Song, 2009], compressed sensing [Bai et al., 2015], various constraints [Benjamini and Basser, 2016], Bayesian estimation [McGivney et al., 2018], or deep learning [Pirk et al., 2020] to improve spectral resolution.

In general, the presence of the fixative and the reduced temperature (room temperature vs. body temperature) decreases the diffusivities in fixed tissues compared to those observed in the live human brain [Dyrby et al., 2011]. It is important to note that if we scale all diffusivities by any factor, say 3, and the b-values used in our experiment by its inverse, i.e., 1/3, all signal attenuations, *e*^-*bD*^, remain unchanged. Consequently, the Monte Carlo simulations with different SNR levels obtained using fixed brain diffusivities and this study’s experimental design with *b_max_* = 10,000*s*/*mm*^2^ also accurately describe an experiment in which all tissue diffusivities are scaled by a factor of 3 simulating *in vivo* conditions and all b-values are scaled by a factor of 1/3, i.e., *b_max_* = 3, 333*s*/*mm*^2^, simulating clinical scan parameters.

### Comparison of dMRI and histological sections

Figure 5 shows a multi-scale side-by-side comparison of a coronal section stained with SMI-32 and the corresponding dMRI data in a representative region of the dorsal premotor cortex. At the macroscopic scale (Fig. 5A,B) we can clearly see that the dominant diffusion direction in the FOD direction-encoded colored (DEC) image [Dhollander et al., 2015] (Fig. 5B) varies continuously along the cortical ribbon and remains perpendicular to the cortical surface. At the mesoscopic scale (Fig. 5C,D) the curvature of the cortex becomes less prominent and the tissue architecture reveals radially oriented neurofilaments in pyramidal neurons with a staining intensity that varies in a laminar pattern reflecting distinct cortical layers. The FODs measured with dMRI in the same region (Fig. 5C) show a good alignment of water diffusion with the dominant orientation of the local microstructure at the scale of hundreds of micrometers. A careful visual inspection of the SMI-32 section at high magnification (Fig. 5E) reveals the presence of cell processes oriented radially and tangentially with respect to the cortical surface. The contribution of tangential processes contributes to the slight differences in SMI-32 staining intensities across cortical layers. At this scale, the grid-like cortical architecture is clearly observable in the orthogonal orientations of the FOD peaks which vary continuously and coherently across multiple voxels (Fig. 5F). These observations confirm similar results from numerous high-resolution dMRI studies and suggest that cortical diffusion processes are locally oriented along orthogonal reference frames that match the tissue architecture and do not change significantly at the scale of a few hundred micrometers, providing a strong justification for describing diffusion processes at smaller length scales with the same fixed locally orthogonal reference frame.

**Figure 5.**
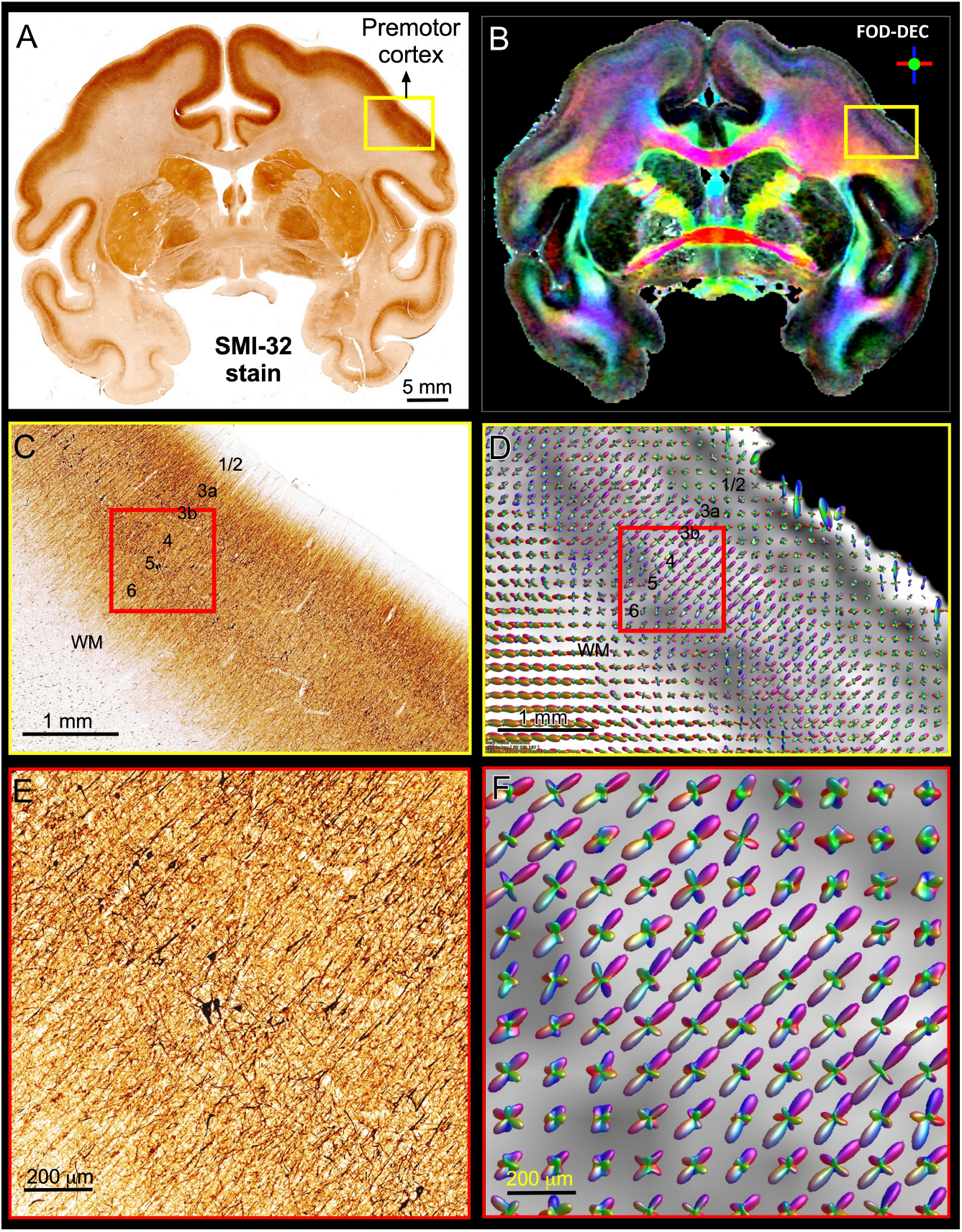
Views of the brain anatomy at the macroscopic scale in a coronal tissue section stained with SMI-32 (**A**) and the FOD-DEC image in a matched MRI slice (**B**) showing the dependence of the principal diffusion direction on the cortical folding geometry. **C** and **D**: Enlarged views of the mesoscopic scale of the histological image and FOD glyphs corresponding to the yellow outlines in **A** and **B**, respectively. The cortical architecture shows a laminar pattern of radially coherent cell processes with different densities (labeled cortical layers). **E** and **F**: Enlarged views of the histological image and FOD glyphs corresponding to the red outline in **C** and **D**. The locally coherent alignment of FOD peaks (**F**) matches the microstructural tissue architecture comprising radial and tangential cell processes (**E**).

### Cortical architectonic features revealed with cDTD MRI

The SNR was estimated as the non-diffusion attenuated magnitude signal averaged in a region-of-interest (ROI) divided by the noise standard deviation measured in an ROI outside the brain using the raw magnitude signals (before post-processing). The cortical SNR varied between 50 and 120. Several imaging artifacts may contribute to an underestimation of the SNR, including:

1. ghosting/aliasing artifacts induced by the vibration of gradient coils (potentially leading to noise overestimation)
2. inaccurate calibration of the transmit and receive gains causing a non-zero background in the reconstructed images (potentially leading to noise overestimation), and
3. spatial inhomogeneities in the B1 sensitivity (potentially leading to tissue signal underestimation)

Our preliminary results of imaging 2D cDTDs in cortical GM reveal diffusion processes with distinct joint radial and tangential diffusivities and different specificities across cortical domains and layers. In Fig. 6, the spectral component images on the diagonal line λ_*r*_ = λ_*t*_ represent isotropic diffusion processes, while those below and above this line quantify anisotropic processes that can be described using prolate and oblate diffusion tensors, respectively. Comparing the maps of the 1D marginal distributions of λ_*r*_ (Fig. 6, left column) and λ_*t*_ (Fig. 6, top row) we found that the spectra of radial diffusivities in tissue microenvironments provides slightly better sensitivity to cortical layers than those of tangential diffusivities. Fig. 6B quantitatively maps the concentrations of eight distinct microscopic diffusion processes and were computed by integrating the 2D cDTDs over spectral domains (Fig. 6A, color-coded outlines) defined empirically based on spatial correlations of spectral components. The resulting signal component maps show high specificity to various cortical layers and were in good agreement with the diffusion orientational features observed in the FOD maps (Fig. 6C). For example, high concentrations of radial microscopic diffusion processes were observed primarily in the mid-cortical layers (Fig. 6B, Components 1 and 7) and in subcortical WM (Fig. 6B, Component 2), while high concentrations of more isotropic and tangential microscopic diffusion processes were observed primarily in the superficial and deep cortical layers (Fig. 6B, Components 5 and 8). The spatial distribution of Component 3 (Fig. 6B) in layer 3 and part of layers 5 and 6 matched with the distribution of non-pyramidal neurons in the parvalbumin stained section (not shown in Fig. 6). Meanwhile, the dense and patchy distribution of Component 6 (Fig. 6B) localized mainly in layer 5 corresponded to the intensely stained pyramidal neurons in this layer in AchE- (not shown in Fig. 6) and SMI-32-stained sections (Fig. 6D).

**Figure 6.**
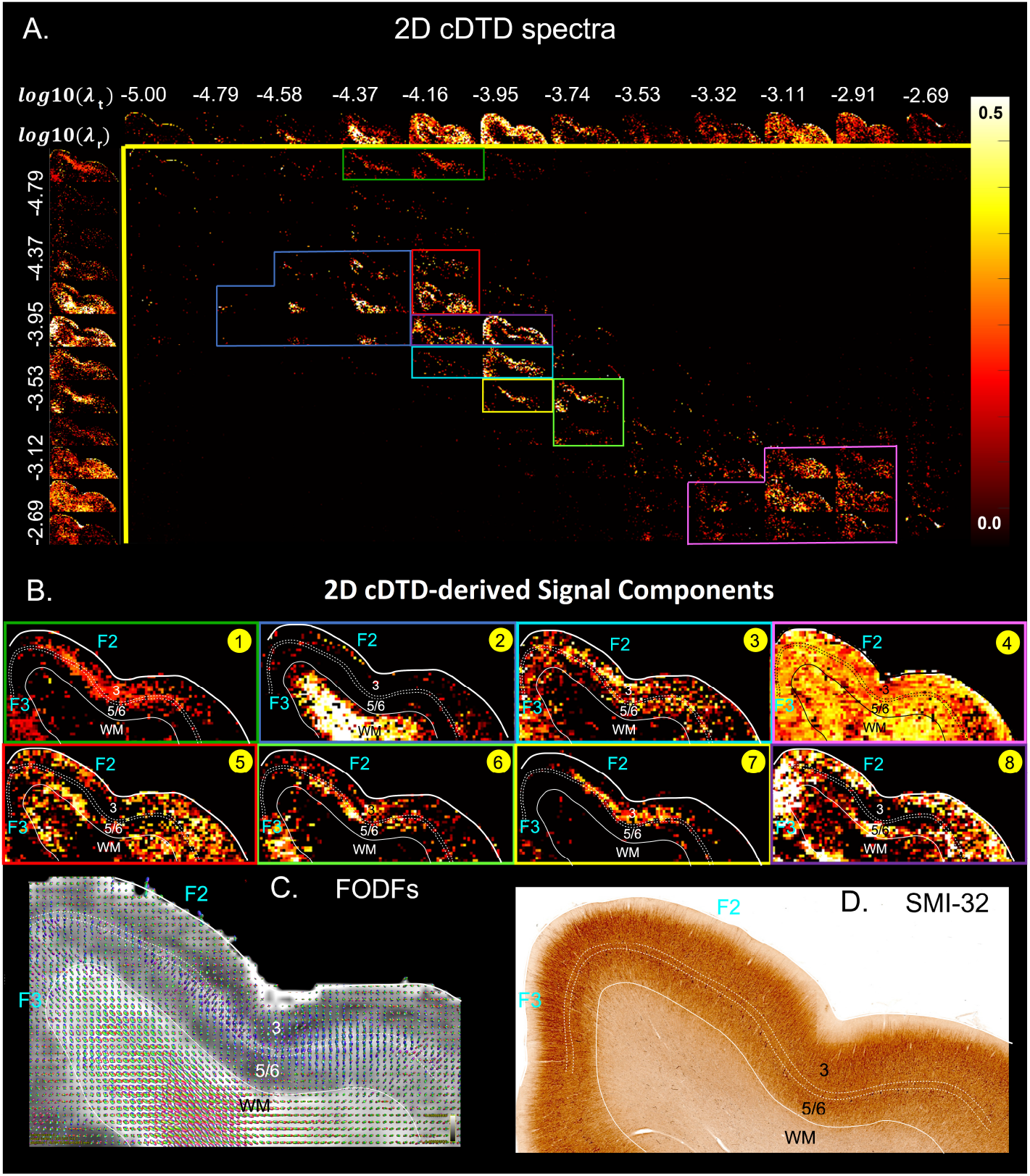
**A.** Spectral component maps of normalized 2D correlation spectra of radial and tangential diffusivities in a section of the cortex from **Fig 5A**. **Top row**: Spectral component maps of the normalized 1D marginal distribution of tangential diffusivity, λ_*t*_; **Left column**: Spectral component maps of the normalized 1D marginal distribution of radial diffusivity, λ_*r*_. **B.** Tissue component maps derived by integrating the 2D cDTD spectral components over empirically defined spectral regions of interest delineated with different colors show good specificity to cortical layers. **C.** Corresponding FODs. **D.** Corresponding SMI-32 stained section.

### Shape-size correlation spectra derived from the cDTD distributions

The 2D *μFA* – *MD* correlation spectral amplitude maps in Fig. 7 provide a tally of the shape-size characteristics of the microscopic diffusion tensors of the DTD as a new means to characterize tissue microstructure. The largest concentrations of isotropic microscopic diffusion processes (*μFA* < 0.18) were observed in the upper cortical layers, and to a lesser extent, in layer 5. The most anisotropic diffusion processes (*μFA* > 0.35) were localized in the mid cortical layers and in the subcortical white matter. The signal in subcortical WM voxels spanned a large range of *μFA* values, potentially reflecting diffusion processes with a larger intravoxel orientational variance (e.g., bending/crossing WM fibers) that may be inadequately described by the cDTDs. The 1D marginal distributions of both the microscopic fractional anisotropies (Fig. 7A, top row) and mean diffusivities (Fig. 7A, left column) derived from the *μFA* – *MD* spectra show layer-specific motifs that allow us to distinguish between superficial, mid, and deep cortical layers. Spectra of MD values in microscopic water pools show the highest concentration of low MD processes in WM (Fig. 7A, component 3), and a mixture of diffusion processes with low and high water mobilities in the mid-cortical layers, potentially indicating important differences in cellularity between these layers. Meanwhile, spectra of *μFA* values revealed predominantly anisotropic diffusion processes in the mid-cortical layers and more isotropic diffusion processes in the superficial and deep layers. Fig. 7B quantifies the spatial distributions and concentrations of five distinct microscopic diffusion components obtained by integrating the 2D *μFA – MD* correlation spectra over empirically defined spectral domains (Fig. 7B, color-coded outlines). In Fig. 7B, Components 1, 3, and 4 are specific to the midcortical layers, while Components 2 and 5 are localized almost exclusively in the superficial/deep cortical layers and in subcortical WM, respectively. Component 3 in the *μFA – MD* maps (Fig. 7B), shows very high *μFA* and likely corresponds in part to the signal from Component 1 in the λ_*r*_ – λ_*t*_ maps (Fig. 6B) with a small λ_*r*_ and large λ_*t*_. It appears to suggest the presence of a small concentration of highly anisotropic oblate microscopic diffusion tensors.

**Figure 7.**
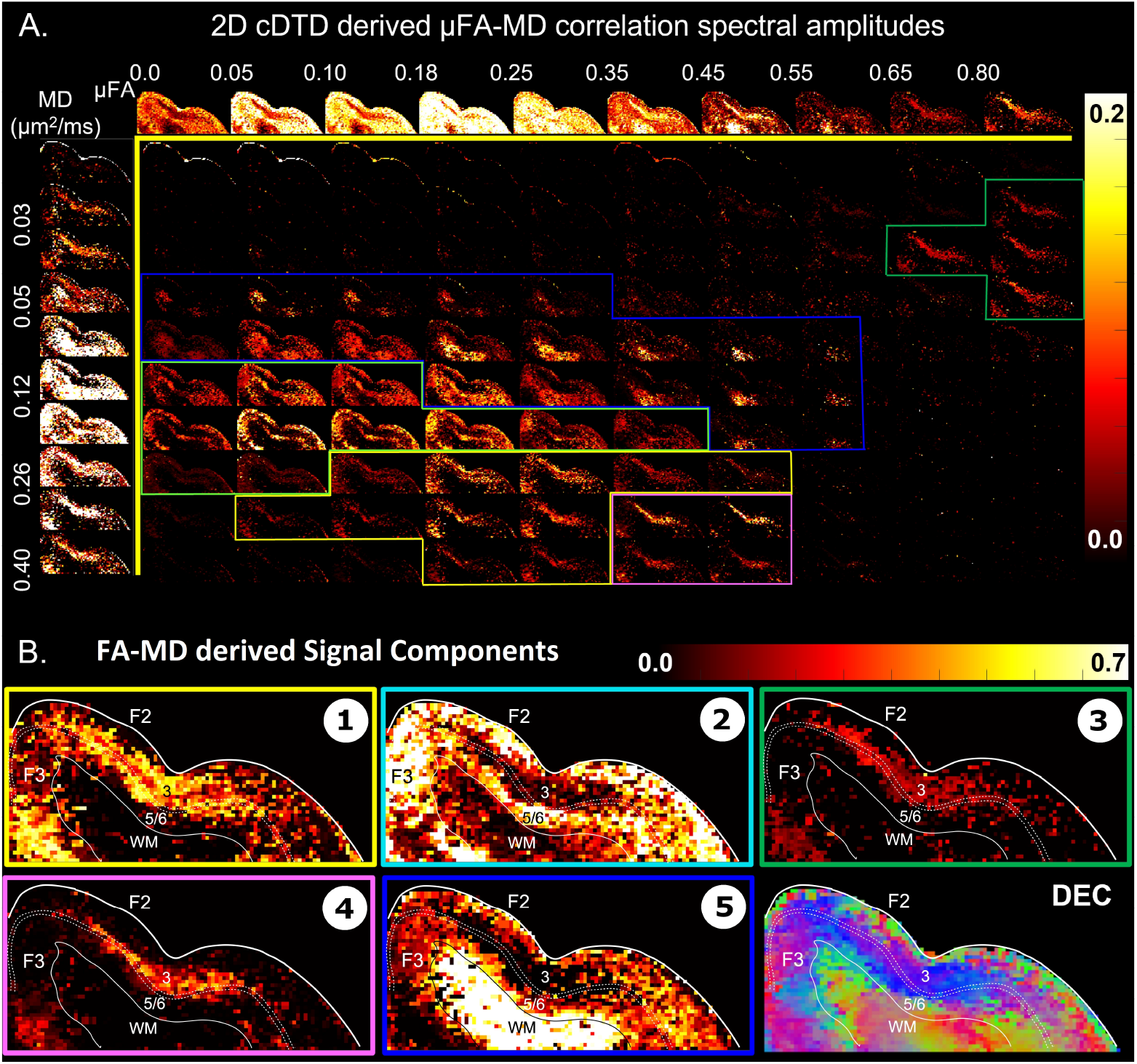
**A.** Spectral amplitude maps of normalized 2D *μFA – MD* correlation spectra in the section of the cortex from **Fig. 6**. **Top row**: Spectral component maps of the normalized 1D marginal distribution of microscopic fractional anisotropy, *μFA*; **Left column**: Spectral component maps of the normalized 1D marginal distribution of the microscopic diffusion tensor mean diffusivities. **B.** Tissue component maps derived by integrating the 2D *μFA – MD* distributions over empirically defined spectral regions reveal strong contrast in the mid-cortical areas.

It is likely that this component reflects restricted water diffusion within tangentially oriented tissue and cell processes (e.g., neurites, neurofilaments) which are powder-averaged within the plane of the mid-cortical layers (Fig. 5E). In this case, the restricted tangential diffusion processes cannot be accurately modeled using tensors (e.g., a powder-average of prolate tensors) and the tangential diffusivities derived with DTD MRI, in general, do not accurately reflect the water diffusivities in different pools (e.g., inside or outside the dendrites). Nevertheless, even if the cDTD-derived diffusivity and anisotropies spectral components may not be quantitative (i.e., biased), they could still provide important clinical information about the density of tangentially oriented neurites or the transverse tortuosity of the extracellular space.

### Potential sources of errors

The accuracy of the measured cDTD spectra depends on several experimental factors such as the number of measurements, the diffusion gradient directions, b-values, as well as SNR. During the voxel-wise cDTD reconstruction, the dMRI signals are decomposed along the axes of the local frame of reference. Consequently, for the same diffusion encoding (i.e., same DWI) the effective diffusion weightings (Eq. 6) of the radial and tangential diffusivities, *b* cos^2^ *ϕ*_**g**_ and *b* sin^2^ *ϕ*_**g**_, respectively, may differ from voxel to voxel. To prevent biases due to the orientations of the local microstructure in the reconstructed cDTD maps it is important that the diffusion encodings uniformly sample the unit sphere for each b-value and across b-values.

Two additional potential sources of errors in the spatial-spectral mapping of microscopic diffusion processes with CORTECS MRI in this study may arise from 1. inaccuracies in estimating the DTI-derived reference frame, and 2. inconsistencies between the axes of the DTI-derived reference frames across neighboring voxels due to the sorting bias of the diffusion tensor eigenvalue decomposition [Pierpaoli and Basser, 1996]. Both sources of errors become more prominent when the dMRI voxel signal is more isotropic. If the signal is isotropic in 3D, the principal diffusion axes are poorly-defined and the estimated diffusion reference frames may be inconsistent across adjacent voxels.

In cortical tissues, the DTI and, more generally, the dMRI signals are radially symmetric even at high spatial resolutions and high b-values. As a result, it is difficult to uniquely define orthogonal principal diffusion axes within the tangential orientation. Instead, we can use a more economical characterization of the microscopic diffusion processes using a distribution of axisymmetric tensors. The resulting 2D cDTDs are completely defined by the correlation spectrum of radial and tangential diffusivities and the dominant diffusion direction (i.e., radial orientation), which can be reliably estimated in the cortex. The Diffusion-Encoded Color (DEC) map in Fig. 5 shows a continuously varying radial diffusion orientation along the cortical ribbon. Despite variations in diffusion anisotropy across cortical layers the principal axis of diffusion corresponding to the largest DTI eigenvalue, *ϵ*_**1**_*ϵ*_**1**_^*T*^, can be reliably estimated throughout the cortex and is consistently oriented normal to the cortical surface. Moreover, this orientation matches that of the largest FOD peak in each corresponding voxel. The side peaks of the FODs are consistently oriented in the tangential plane perpendicular to the radial direction, supporting the orthogonal alignment of diffusion processes, in good agreement with findings from previous high-resolution cortical dMRI studies [Aggarwal et al., 2015, Kleinnijenhuis et al., 2013, Leuze et al., 2014].

However, more generally, when DTI data is acquired with lower spatial resolution, low FA values in the cortex can bias the measurement of the radial direction that determines the 2D cDTD reference frame in each voxel. In this situation, it may be possible to use higher b-values (or longer diffusion times) to improve the sensitivity to the orientational features of the dMRI signal, and/or to estimate the voxel reference frame more reliably from the directions of the largest FOD peaks. Alternatively, one could derive a cortical reference frame from the curvature of the cortex measured using a structural scan with good GM-WM contrast as a proxy for the diffusion reference frame [Avram et al., 2020] or use spline interpolation of the diffusion tensor field [Pajevic et al., 2002] in low FA voxels, to derive a continuously varying reference frame that is consistent throughout the cortex.

## 1 Discussion

The CORTECS framework greatly simplifies the data acquisition and spectral reconstruction requirements for high-resolution DTD MRI and subsumes many previously proposed diffusion tensor models. It provides a practical and feasible approach to non-parametric quantitation of microstructural heterogeneity in healthy and diseased tissues. At its core, the framework relies on the observation that, in tissues with consistent well-defined architecture, such as the cortex, as we increase the spatial resolution from the scale of a conventional dMRI voxel (≈ 2*mm*) relative to the radius of curvature of the underlying anatomy, the intravoxel angular dispersion of diffusion processes decreases. At the mesoscopic scale of a few hundred micrometers diffusion processes in distinct tissue microenvironments, e.g., associated with myelin, intra-, extra-axonal water, remain largely coincident along the axes of a common reference frame determined by the local tissue architecture. At this length scale, the intravoxel angular dispersion due to cortical folding is significantly reduced and differences between subvoxel (microscopic) diffusion processes are primarily characterized by their principal diffusivities. Correlations between principal diffusivities explain most of the microscopic diffusion heterogeneity. They determine the anisotropies and mean diffusivities of the microscopic diffusion tensors, i.e., the shapes and sizes of their diffusion ellipsoids, rather than their relative orientations, allowing us to constrain the DTD reconstruction.

### The persistence of the principal diffusion orientations for various signal weightings

The basis of constraining cortical diffusion processes to be oriented along local orthogonal directions in neural tissue has many lines of support. Direct observations of cortical cyto- and myelo-architectonic features with optical and 3D electron microscopy reveal dominant radial and tangential orientations. Meanwhile, histological validation studies using high spatial and angular resolution dMRI with a range of mesoscopic spatial resolutions have repeatedly shown that in neural tissues the preferential diffusion directions align with the dominant orientation of the underlying microstructure. Moreover, results from numerous high-resolution dMRI studies suggest that when the relative signal contributions (weightings) from specific water pools are altered using different signal preparations the principal axes of the diffusion tensors and the orientations of the dominant FOD peaks in the voxel do not change [Assaf, 2019]. Concretely, the dominant diffusion orientations do not change significantly in experiments with a wide range of echo times (T2-weightings) [Avram, 2011, Avram et al., 2012], repetition times, inversion times (T1-weightings) [Assaf, 2019], b-values (diffusivity weightings) and diffusion times (chemical exchange and restriction weightings). Furthermore, in vivo experiments combining diffusion MRI and magnetization transfer (MT) preparation indicate that in white matter fibers the principal diffusion directions of myelin water and non-myelin water pools are coincident [Avram et al., 2010]. Similarly, in vivo diffusion tensor spectroscopy experiments of neuronal-specific metabolites, such as NAA have shown that diffusion processes in intra- and extracellular water pools are also aligned with the diffusion reference frame of the voxel [Ronen et al., 2013]. The persistence of the reference frame under various signal preparations suggests that the intravoxel orientational heterogeneity is dominated by the curvature of the macroscopic anatomy (e.g., cortical folding, fanning/bending WM pathways), and that water diffusion in specific microenvironments of neural tissues can be described adequately with a singular reference frame defined by the mesoscopic architecture. Finally, constraining subvoxel cortical diffusion tensor processes to the local reference frame of the mesoscopic voxel may also be justified with arguments from developmental biology.

### Orthogonal reference frames in neurodevelopment

During morphogenesis, diffusion-reaction processes can establish orthogonal concentration gradients [Gregor et al., 2005, Turing, 1952] to support the efficient transport of macromolecules such as growth and inhibitory factors. It is believed that in early embryogenesis this mechanism [Gregor et al., 2005, Lefèvre and Mangin, 2010] leads to the formation of the principal axes of embryonic development: rostro-caudal, medio-lateral, and dorso-ventral [Kingsbury, 1920]. Similarly, during early brain development diffusion-reaction processes at the microscopic scale, e.g., ≈ 10 – 50μm, likely guide the growth of elongated cellular and sub-cellular structures, such as neurofilaments, axons and dendrites, which in turn, provide a scaffold for the diffusive migration and active transport of macromolecules over longer distances. The progressive elaboration of the orthogonal reference frame provides a plausible explanation for the architecture of cortical columns, laminae, and capillaries, at the mesoscopic scales of ≈ 100 – 500*μm*. Diffusion MRI studies in the late stages of fetal neurodevelopment and newborns have shown a decrease in the radial coherence of diffusion processes [Dudink et al., 2015, Khan et al., 2019, McKinstry et al., 2002, Takahashi et al., 2011, Vasung et al., 2010].

More generally, several theories of brain development [Chen et al., 2013, Lefèvre and Mangin, 2010, Van Essen, 1997, Wedeen et al., 2012] suggest to different extents, that similar locally orthogonal reference frames may be observed in WM at high spatial resolution. The intravoxel angular dispersion in WM voxels depends on the curvature of the fiber pathways (e.g., due to bending and fanning) as well as the presence of fiber crossings. The radii of curvature due to bending (e.g., corpus callosum) or fanning (e.g., corticospinal tract) in WM pathways are typically larger than those of the cortical folding geometry (e.g., sulci and gyri), even for short-range U-fibers. Consequently, at the mesoscopic spatial resolutions required for CORTECS MRI, the residual intravoxel orientational variation of diffusion processes in WM is due primarily to the crossing angles of subvoxel fiber populations. CORTECS MRI may be applicable in regions containing a single homogeneous WM pathway (i.e., no crossings), such as the corpus callosum, but not in most WM voxels that contain fiber populations that do not cross at orthogonal orientations. Nevertheless, the framework could provide an independent method to test the hypothesized local orthogonality [Tax et al., 2016, 2017] at various spatial resolutions.

### The dimensionality reduction of cDTDs

Current approaches for imaging DTDs and/or their features require SDE and MDE measurements and include parametric models using SDE [Jian et al., 2007] and combinations of SDE and MDE measurements [Henriques et al., 2020, Magdoom et al., 2021, Szczepankiewicz et al., 2016, Westin et al., 2016] as well as non-parametric methods [Topgaard, 2017]. Parametric DTD models approximate the solution using analytical functions such as a Wishart distribution [Jian et al., 2007] or a constrained normal tensor-variate distribution [Magdoom et al., 2021]. While such analytical approximations can estimate DTDs from fewer measurements and lower SNR levels, they drastically limit the space of admissible DTDs to those described by a handful of degrees of freedom (i.e., parameters or coefficients). The reconstructed DTDs may provide biased assessments in voxels affected by partial volume contributions from tissues with very different diffusion properties and may not accurately capture the range of unknown tissue alterations that occur in disease. Non-parametric or spectroscopic DTD reconstruction methods [Topgaard, 2017] can describe an arbitrary range of tissue compositions but, due to the large spectral dimensionality of the problem, require many MDE DWIs with high SNR and computationally intensive statistical reconstruction methods to enforce positive definiteness of the solution.

For a general, unconstrained non-parametric DTD, the microscopic diffusion tensors can have arbitrary orientations (Eq. 2). Consequently, the 6-dimensional random variable of the DTD must support both positive and negative off-diagonal tensor elements and cannot be analyzed with conventional ILT methods. To overcome this limitation, the DTD reconstruction requires computationally intensive statistical methods [Magdoom et al., 2021, Topgaard, 2017] to enforce positive definiteness constraints that ensure the physicality of the microscopic diffusion tensors. Alternatively, if we describe the DTD using the principal diffusivities, λ_1_, λ_2_, λ_3_ and the three Euler angles *ϕ*, *ψ, θ*, which define the orientations of the orthonormal directions *ϵ*_1_, *ϵ*_2_, *ϵ*_3_ in Eq. 1, then *ϕ, ψ, θ* create a trigonometric dependence in the signal equation. The key insight of the CORTECS MRI framework is that in tissues with well-defined, orthogonal architectures, sampling the spatial dimensions more densely, i.e., increasing the spatial resolution, reduces the intravoxel angular dispersion. This allows us to restrict the 3 degrees of freedom that determine the orientations of the tensor random variable, i.e. the three Euler angles, and thus reduce the domain of the DTD to the orthogonal non-negative 3D space of principal diffusivity random variables that guarantees positive definiteness and can be solved with a conventional ILT reconstruction techniques. This trade-off between spatial resolution and spectral dimensionality has several important implications for the clinical translation of non-parametric DTD MRI.

### Data acquisition requirements for CORTECS MRI

In general, the SNR requirements for multidimensional spectral (i.e., non-parametric) reconstruction algorithms scale exponentially with the dimensionality of the problem. For a 2D spectral reconstruction, an SNR of 100 allows us to measure signal attenuations by a factor of 10 along two independent spectral dimensions. Meanwhile, to achieve the same effective dynamic range per dimension for a 4D spectral reconstruction, we need an SNR of 10,000. While such nominal SNR levels may be achievable on clinical scanners by using sufficiently large voxel sizes, the integrity of the data acquired *in vivo* may be corrupted [Avram et al., 2019, 2021] by:

1. imaging artifacts such as ghosting/aliasing, eddy current induced distortions, or Gibbs ringing, which typically represent ≈ 1 – 2% of the tissue signal; and
2. partial volume inconsistencies across DWIs due to subject and physiological motion (e.g., blood flow, pulsations, etc.).

In routine clinical MRI scans, e.g., T1W, T2W, DTI, typical SNR levels are between 20-50, and these signal artifacts on the order of ≈ 2% are barely visible. However, for an in vivo SNR = 10,000, these signal instabilities produce an artifact-to-noise ratio of 200, potentially biasing the estimation of non-parametric DTDs in high dimensional spaces (e.g., 4D or 6D) and rendering them unsuitable for clinical translation.

On the other hand, CORTECS MRI measures 3D or 2D correlation spectra using efficient diffusion preparations (SDEs), fewer measurement encodings (data points), and SNR levels that may be achieved for ultra-high resolution in vivo dMRI in the near future. Advances in various technologies including the design of high-field MRI scanners [Feinberg et al., 2021], high-performance gradient coils [Feinberg et al., 2021, Foo et al., 2020, Huang et al., 2021], high-density RF coil arrays [Hendriks et al., 2019, Keil et al., 2013], as well as efficient high-resolution dMRI pulse sequences [Avram et al., 2014b, Feinberg et al., 2010, Setsompop et al., 2018], image acquisition and reconstruction strategies [Feinberg et al., 2010, Setsompop et al., 2018], and experimental protocols [Avram et al., 2018, 2019, Nilsson et al., 2020] can be integrated synergistically in state-of-the-art MRI systems [Feinberg et al., 2021, Foo et al., 2020, Huang et al., 2021] to achieve the spatial resolution, scanning efficiency, and diffusion sensitizations required for in vivo CORTECS MRI.

In our experiment the acquisition of each high-resolution DWI volume required 52minutes. This relatively long duration scan duration is due to the use of:

1. a large imaging matrix of 375×320×230 needed for whole-brain coverage at 200*μm* resolution, and
2. 3D diffusion spin echo EPI sequence with segmented k-space acquisition and a relatively long TR of 650ms.

The TR was chosen so as to minimize gradient heating (i.e., limit the gradient duty cycle), and included a 150ms duration for excitation, diffusion preparation, and EPI readout, and a 500ms idle duration. For clinical imaging, both factors can be significantly reduced. Firstly, using a multi-slice spin-echo diffusion EPI sequence with multiband capabilities one could acquire each DWI volume efficiently (negligible idle duration) in a single TR of 5-10s, albeit at a lower SNR. Secondly, it is important to point out that the requirement for high spatial resolution in CORTECS MRI does not necessarily require a prohibitively long scan duration. Unlike dMRI fiber tractography, CORTECS dMRI does not require whole-brain data. Using outer-volume suppression, reduced FOV, or ZOOM EPI one could significantly reduce the imaging matrix size and scan duration while still maintaining the required spatial resolution for in vivo scans with human subjects. On the other hand, the scan duration requirement of conventional non-parametric DTD methods is inherently limited by the very large number of encodings needed to sample the high-dimensional space exhaustively, even when scanning with a reduced FOV.

### Spatial resolution requirement in CORTECS MRI

The major drawback of CORTECS MRI compared to conventional (unconstrained) nonparametric DTD methods is the prerequisite of sufficiently high spatial resolution (≈ 400*μm*). The spatial resolution at which we can adopt a common reference frame for all subvoxel diffusion tensors depends on the cortical folding geometry and may vary across the brain. A useful quantity to characterize the validity of this assumption is the dimensionless ratio between the voxel length, *x*, and the minimum radius of curvature of the macroscopic anatomy (e.g., cortical folding, or bending/fanning of WM fibers), *R*. If this ratio is sufficiently small 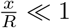, then we can ignore orientational variations of subvoxel diffusion processes (Fig. 1). For example given a voxel size of *x* = 0.2*mm* the expected maximum intravoxel angular variance of the microstructural reference frame due to the continuously varying cortical folding geometry is ±1.9° for *R* = 5*mm* and ±4.9° for *R* = 2*mm*. This angular variation is smaller than even the most ambitious estimates of angular resolution limits in diffusion MRI fiber tractography and is unlikely to bias the estimated spectra. HARDI experiments using well-calibrated diffusion phantoms with overlapping, highly anisotropic coherent structures oriented at different angles cannot typically resolve diffusion processes due to fibers crossing at angles < 10°, even when a large number of gradient orientations with large b-values and high SNR levels are used in microimaging or clinical scanners [Guise et al., 2016, Perrin et al., 2005]. This angular resolution limit provides a good benchmark for the ability to accurately resolve the orientations of subvoxel diffusion tensor processes with conventional 6D nonparametric DTD MRI methods.

The high spatial resolution requirement in CORTECS MRI can lead to significantly longer acquisition time per volume (i.e., per diffusion encoding), when compared to conventional (unconstrained) nonparametric DTD MRI methods. These methods require large imaging voxel volumes to achieve the very high SNR and signal dynamic range needed for 6D or 4D DTD reconstructions and can be affected by signal artifacts. Moreover, these methods also require a large number of joint (multidimensional) econdings to comprehensively sample the high-dimensional parameter space, thereby offsetting potential savings in the total scan duration that may be gained by imaging a smaller matrix size (i.e., larger voxels), when compared to CORTECS MRI. Most importantly, however, the 6D DTDs measured in voxels of ≈ 3*mm* do not provide any information about the relative spatial distribution of subvoxel diffusion tensors, i.e., at length scales smaller than ≈ 3*mm*. Due to its high spatial resolution requirement, CORTECS MRI explicitly measures the relative spatial distributions (and relative orientations!) of diffusion tensor processes at much finer length scales, e.g., down to 200μm in our study, providing significantly more information. Compared to conventional DTD methods, this higher spatial resolution in CORTECS MRI provides more accurate localization and improved sensitivity in the detection of subtle pathological tissue changes, for example in the early stages of neurodegeneration.

### Potential for quantifying diffusion time dependence

All DTD MRI methods assume that the voxel can be viewed as an ensemble of non-exchanging Gaussian (i.e., freely diffusing) subvoxel water pools within which the diffusive motions of spins are described with tensors whose corresponding ellipsoids have different sizes, shapes, and orientations. In biological tissues, cellular and subcellular structures can present microscopic restrictions and hindrances producing a time-dependent (non-Gaussian) diffusion in certain water pools. To address this limitation, the MDE-based DTD frameworks [Topgaard, 2017], can be extended to include diffusion time dependence [Lundell et al., 2019], and/or analyzed using parametric models [Henriques et al., 2020]. The characteristics of time-dependent DTDs can yield important tissue microstructural information about the distribution of compartment shapes and sizes [Henriques et al., 2020, Lundell et al., 2019] that classical MDE experiments sought to measure [Avram et al., 2013b, Benjamini et al., 2016, Koch and Finsterbusch, 2008, Komlosh et al., 2018]. However, it can be troublesome to incorporate the dependence of diffusion processes on the time-varying diffusion gradient waveforms into the signal equation, even for MDE preparations with well-defined diffusion time parameters such as those using double pulsed field gradients [Avram et al., 2013b, Mitra, 1995], or rotating field gradients [Avram et al., 2014a]. Conversely, the diffusion time dependence of SDE measurements can provide similar information to MDE measurements [Jespersen, 2012] and is described by a well-defined parameter △, the separation between the start times of the two diffusion gradient pulses. Moreover, since the voxel reference frame does not change significantly with diffusion time [Assaf, 2019], we can directly extend the CORTECS framework to map time-dependent cDTDs by repeating the experiment with multiple diffusion times. Imaging correlation spectra of diffusion-time-dependent principal diffusivities in microscopic water pools may provide important pathophysiological information about microscopic restrictions, chemical exchange, and water transport [Nilsson et al., 2013].

### Relation to other dMRI methods

The non-parametric cDTD signal representation can be viewed as a multi-tensor generalization of high-resolution DTI. It subsumes many parametric tissue diffusion models for WM [Stanisz et al., 1997] and GM [Avram et al., 2020, Mulkern et al., 1999] and enables their cross-validation. It can inform the design of more efficient dMRI experiments using SDEs and MDEs to measure parametric DTDs and tensor mixture models for specific clinical applications. Moreover, it provides an independent method for deriving DTD-related quantities, such as the non-parametric distribution of subvoxel MD values which can be measured efficiently in a 6 min clinical scan [Avram et al., 2019]. In this way, the proposed framework may help test the validity of various DTD methods and guide their development towards achieving higher spatial resolution and greater biological specificity.

The ability to quantify tissue properties non-parametrically is crucial to our understanding of disease progression, tissue regeneration, and neurodevelopment. By quantifying subvoxel DTDs non-parametrically we can identify the most prominent spectral features such as the shapes and peaks or multimodal clusters associated with specific pathophysiological changes. Once we learn these spectral signatures, we can model the CORTECS-derived 2D or 3D cDTDs using analytical functions determined by only a few parameters. Disease-specific parametric cDTD could be reconstructed swiftly and efficiently from data acquired with lower SNR and a smaller number of encodings.

### Further improvements in biological specificity

The correlation spectrum of principal diffusivities may reveal signal contributions from specific tissue components, such as intra-axonal, extracellular, or myelin water whose diffusion tensors may be coincident and are therefore difficult to disentangle based on orientational diffusion characteristics such as FODs derived from HARDI data. A further improvement in biological specificity may be achieved by integrating the cDTD measurements with multidimensional relaxation MRI methods [Benjamini and Basser, 2017, Kim et al., 2017] which measure the net voxel signal as a superposition of contributions from subvoxel water pools with different joint T1-, T2- and diffusion properties. However, with the addition of new dimensions for contrast encoding, most implementations of diffusion-relaxation correlation MRI on clinical scanners require larger datasets, higher SNR levels as well as the use of sophisticated pulse sequences and algorithms to reconstruct five-dimensional [Reymbaut et al., 2021] or six-dimensional [de Almeida Martins et al., 2021] correlation spectra. We have recently proposed a more practical two-dimensional diffusion-relaxation MRI method for efficiently mapping T1-MD correlation spectra using isotropic diffusion encoded (IDE) DWIs [Avram et al., 2021]. Similarly, the CORTECS framework adds the minimum number of dimensions (principal diffusivities) needed to efficiently combine T1- or T2-relaxation with diffusion tensor spectroscopic imaging.

### Potential applications to neuroscience and neuroradiology

Mapping water pools in specific cortical microenvironments based on their diffusion tensor properties quantitatively and efficiently could have numerous applications in neuroradiology and neuroscience. It may improve the diagnosis of neurodevelopmental disorders and allow us to specifically disentangle contributions from increased dendritic arborization and reductions in radial glial fibers to the cortical microstructural changes observed in newborns. In addition, it may provide biomarkers for early detection of cortical microstructural changes occurring in epilepsy [Lampinen et al., 2020], cancer [Szczepankiewicz et al., 2016], traumatic brain injury [Komlosh et al., 2018], stroke [Alves et al., 2022], or multiple sclerosis [He et al., 2021]. Mapping correlations between cortical diffusion processes with CORTECS MRI could quantify specific cellular/tissue components providing new parameters for automatic cortical parcellation and layer segmentation algorithms. Relating these layer-specific components to input and output signaling in cortical areas could allow us to study intracortical connectivity and gain insight into the directionality of information flows (signaling) in functional networks throughout the connectome [Olman et al., 2012, Uğurbil et al., 2013]. Because it requires only SDE data, CORTECS MRI can be applied retrospectively to analyze existing high-resolution diffusion MRI data sets. Finally, while this study focuses on quantifying diffusion in cortical gray matter, CORTECS MRI may also be applicable to other organized tissues with varying degrees of macroscopic and microscopic diffusion anisotropies such as in white matter, kidney medulla, heart muscle, skeletal muscle, ligaments, tendons, etc.

## 2 Conclusions

This study provides a new framework for empirical and biologically specific analyses of subvoxel diffusion heterogeneity in healthy and diseased brain tissue using conventional high-resolution dMRI. From the non-parametric cDTDs we can derive additional spectral and scalar parameters, such as the joint size-shape distribution of microscopic diffusion tensors. Our preliminary results in the macaque monkey cortex reveal diffusion components that correlate well with distinct architectonic features. CORTECS MRI has the potential to advance the clinical translation of DTD MRI and the optimization for specific applications in clinical and basic sciences. Features of cDTD spectra may help better delineate cortical layers and areas in healthy subjects and may provide new biomarkers for finding subtle cortical abnormalities underlying focal dysplasia in epilepsy, microbleeds in traumatic brain injury, metastatic cancers, etc.

## Acknowledgments

This work was supported by the Intramural Research Program of the *Eunice Kennedy Shriver* National Institute of Child Health and Human Development, “Connectome 2.0: Developing the next generation human MRI scanner for bridging studies of the micro-, meso- and macro-connectome”, NIH BRAIN Initiative 1U01EB026996-01 and the CNRM Neuroradiology-Neuropathology Correlation/Integration Core, 309698-4.01-65310, (CNRM-89-9921). We thank Drs. Michal Komlosh, Cecyl Chern-Chyi Yen, and Frank Ye for assistance with sample preparation and data acquisition, Drs. Paul Taylor and Daniel Glen for helpful discussions and Drs. Bernard Dardzinski and Alexandru Korotcov for providing the RF coil used in this experiment. We thank Tom Pohida and Marcial Garmendia-Cedillos for their help in constructing the 3D brain mold. We also thank Drs. Ted Usdin and Sarah Williams-Avram as well as Drs. Vincent Schram and Ross Lake for help with optical imaging and digitization of microscope slides, and FD Neurotech for histological services provided. Also, we thank Drs. Betsy Murray and Richard Sanders in the Laboratory of Neurophysiology in NIMH for providing the perfusion-fixed monkey brains for our experiments.

The opinions expressed herein are those of the authors and not necessarily representative of those of the Uniformed Services University of the Health Sciences (USUHS), the Department of Defense (DoD), VA, NIH or any other US government agency, or the Henry M. Jackson Foundation.

